# Equivalence of the Erlang Seir Epidemic Model and the Renewal Equation

**DOI:** 10.1101/319574

**Authors:** David Champredon, Jonathan Dushoff, David J.D. Earn

**Affiliations:** Department of Biology, McMaster University, Hamilton, ON, Canada; Department of Mathematics and Statistics, York University, Toronto, ON, Canada.; Department of Biology, M.G. DeGroote Institute for Infectious Disease Research, McMaster University, Hamilton, ON, Canada; Department of Mathematics and Statistics, M.G. DeGroote Institute for Infectious Disease Research, McMaster University, Hamilton, ON, Canada

**Author notes:** **Funding:** This work was funded by the Natural Sciences and Engineering Research Council of Canada and the Canadian Institutes of Health Research.

**Keywords:** epidemic models, renewal equation, differential equations SEIR Erlang, generation interval distribution

## Abstract

Most compartmental epidemic models can be represented using the Euler-Lotka renewal equation (RE). The value of the RE is not widely appreciated in the epidemiological modelling community, perhaps because its equivalence to standard models has not been presented rigorously in non-trivial cases. Here, we provide analytical expressions for the intrinsic generation interval distribution that must be used in the RE in order to yield epidemic dynamics that are identical to those of the susceptible-exposed-infectious-recovered (SEIR) compartmental model with Erlang-distributed latent and infectious periods. This class of models includes the standard (exponentially-distributed) SIR and SEIR models as special cases.

## 1. Background

The renewal equation (RE) was introduced by Leonhard Euler in 1767 [10] in his work on population dynamics and was “rediscovered” in a modern continuous formulation by Lotka in 1907 [20]. Lotka’s formulation is usually expressed as

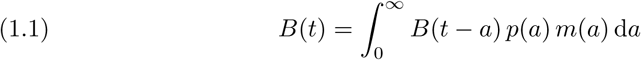

where *B*(*t*) is the number of births at time *t*, *p*(*a*) is the probability of survival to age *a*, and *m*(*a*) is the fertility at age *a*. This equation was derived for demographic studies and has been adapted to model epidemics (for example [9, 21, 22]) by changing the interpretation of the variables: *B*(*t*) represents the number of new infectious individuals at time *t*, *p*(*a*) the probability to be infectious *a* time units after acquiring the disease, and *m*(*a*) the “transmission potential”, that is the average number of secondary infections at “infection age” *a*.

The dynamics of epidemics are more commonly modelled with ordinary differential equations (ODEs), following the seminal work of Kermack and McKendrick in 1927 [17]. This family of models identifies epidemiological states (susceptible, infectious, immune, *etc*.) and considers the flow rates between “compartments” containing individuals in each disease state. A standard example is the “SEIR” model, which distinguishes between a latent state of infection, traditionally labelled *E* for “exposed”, where the infected individual is not yet infectious, and then a state *I* where the infected individual is infectious. When not infected, an individual is either susceptible (*S*) or immune/recovered (*R*). A generalization of this model, which we will call the “Erlang SEIR model”, divides the *E* and *I* stages into *m* and *n* substages, respectively. All *m* latent (respectively *n* infectious) substages are identical. This subdivision is usually viewed as a mathematical trick in order to make latent and infectious period distributions more realistic; the resulting latent and infectious periods have Erlang distributions (Gamma distributions with integer shape parameter) [1, 19, 26, 18].

The renewal and ODE approaches are based on different conceptualizations of dynamics. The renewal approach focuses on cohorts of infectious individuals, and how they spread infection through time, while the ODE approach focuses on counting individuals in different states. The renewal equation is less common than compartmental models in epidemiological applications, probably because the goal when modelling epidemics is often to identify optimal intervention strategies, which is facilitated by clearly distinguishing the various epidemiological states (e.g., susceptible, infectious, immune, vaccinated, quarantined, *etc*.) to act on. However, the simplicity of the renewal equation makes it particularly well adapted to estimate the effective reproductive number from incidence time series [25] and to forecast epidemics [8]. As a notable example, it was used recently by the WHO Ebola Response Team to estimate the reproductive number the Ebola epidemic [27].

Despite their very different formulations, these two models can simulate exactly the same epidemics when the generation-interval distribution *g* derived from the ODE system is used in the renewal equation [11, 28, 5]. However, apart from simple cases with exponential distributions, the generation-interval distribution *g* that links Erlang SEIR models to renewal-equation models has apparently never been explicitly derived. Here, we provide an analytical expression for the intrinsic generation-interval distribution implied by an Erlang SEIR model and show that a renewal equation model using this distribution for *g* yields exactly the same epidemic dynamics as the corresponding compartmental model.

## 2. Methods

In this section, we define the notations and equations for the renewal and Erlang SEIR models. We consider a normalized population (i.e., the total population size is 1) and set the day as the time unit. The computer code for all numerical simulations is provided in Supplementary Material.

### 2.1. The Erlang SEIR model

The Erlang SEIR model, with balanced vital dynamics, is described by a system of *m* + *n* + 1 ODEs,

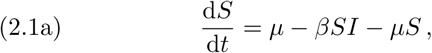

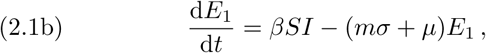

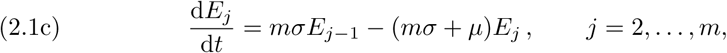

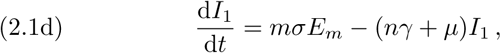

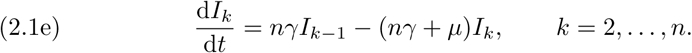

where 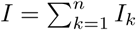. The parameter *β* is the transmission rate, 1/*σ* is the mean latent period (assuming an individual survives latency), 1/(*γ* + *µ*) is the mean duration of infectiousness, and *µ* represents the per capita rates of both birth^1^ and death. To reduce the notational burden, the dependence on time has been omitted (i.e., *S* = *S*(*t*)). Initial conditions are discussed below in §2.4.

The basic reproduction number for the Erlang SEIR model (2.1) is easily derived [24, 13, 18],

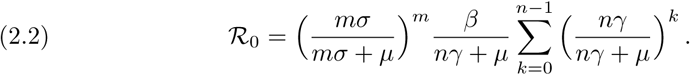

Note that in the absense of vital dynamics (*µ* = 0), this expression reduces to 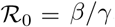.

### 2.2. Intrinsic generation interval distribution via cohort equations

In addition to the ODE system (2.1) describing the number of individuals in different clinical states, we can naturally define another ODE system for the probabilities to be in these different clinical states at a given time after infection. Let *L_j_*(*τ*) be the probability that an individual is alive and in the *j*^th^ latent stage (*E_j_*) at time *τ* after being infected. Similarly, let *F_k_*(*τ*) be the probability that one individual is alive and in the *k*^th^ infectious stage (*I_k_*) at time *τ* after being infected. In other words, we model the proportion in each stage of each infectious cohort.

We have *L*_1_(0) = 1, *L_j_*(0) = 0 for *j* = 2, …, *m* and *F_k_*(0) = 0 for *k* = 1, …, *n*. We construct equations for the *L_j_* and *F_k_* exactly in parallel with the equations for *E_j_* and *I_k_*:

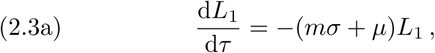

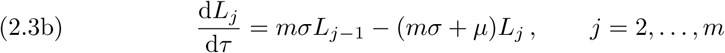

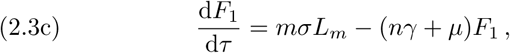

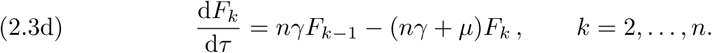

The probability to be infectious at time *τ* after acquiring infection is simply the sum 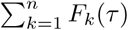 (an individual can only be in one single infectious state at any given time). The intrinsic generation-interval distribution [7] for the Erlang SEIR model, denoted *g*, can then be expressed as

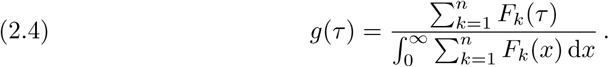

### 2.3. The renewal equation with susceptible depletion

For typical transmissible infections, individuals acquire immunity after recovering and cannot be reinfected (at least for some time). Consequently, the total number of susceptible individuals decreases during an epidemic. In addition, individuals who successfully transmit their infection to others must survive at least until the moment of transmission. Finally, new susceptible individuals are recruited through births, and all individuals have a finite lifespan. To account for these processes of “susceptible depletion”, “survival to transmission” and “vital dynamics” (which are present in the Erlang SEIR model), Lotka’s equation (1.1) must be revised.

As in the ODE model (2.1), we denote by *S*(*t*) the proportion of the population that is susceptible at time *t*. However, unlike the ODE model, our renewal equation will be expressed in terms of *incidence i*(*t*) rather than prevalence *I*(*t*). Incidence is the *rate* at which new infections occur in the population, and corresponds to the flow rate *βSI* from *S* to *E*_1_ in Equation 2.1a. Our renewal equation is

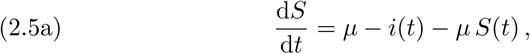

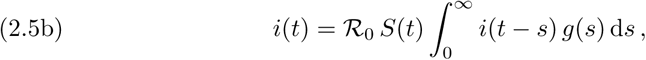

where 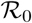 is the basic reproduction number and *g* is the intrinsic generation-interval distribution [7]. The function *g*(*τ*) is the probability that an individual survives and transmits the disease *τ* days after acquiring it. Note that both 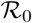 and the distribution *g* implicitly account for deaths of exposed and infectious individuals. This contrasts Equation 2.2, in which 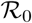 is expressed explicitly (and actually derived) in terms of rate parameters, including the mortality rate *µ*.

### 2.4. Initial conditions

To complete the formulation of the RE model (2.5), we must specify initial conditions. Doing so is not as straightforward as for the ODE model (2.1), for which the initial state is simply an (*m* + *n* + 1)-dimensional vector containing the proportions of the population in each compartment. Instead, in addition to the initial proportion susceptible, *S*(0), for the RE we must specify the incidence at *all* times before *t* = 0, i.e., *i*(*t*) for all *t* ∈ (−∞, 0]. Here, we use the Dirac Delta distribution, *δ*(*t*), to “jump-start” the epidemic at time 0, and write:

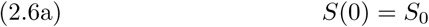

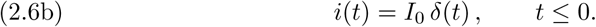

This is equivalent to starting at time 0 with a proportion *I*_0_ in the first infected state (state *I*_1_ if *m* = 0, state *E*_1_ otherwise), and no other infected individuals. The RE (2.5) with these initial conditions (2.6) can be solved numerically in a straightforward manner. Appendix C outlines the algorithm that we have used in our numerical simulations. This approach allows us to simulate efficiently, and to start with any number of susceptible and infected individuals, thus effectively spanning the phase space.

We note that, with more complicated simulations, it would be possible to match not only the number susceptible and the total number infected (as above) but also how the initial prevalence is spread among the *m* + *n* infected classes in the ODE model (2.1), by using an alternative formulation [3] for (2.5b):

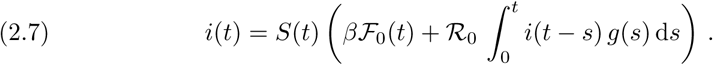

Here, the integral over the generation interval looks back only to time 0 (not time −∞) and the force of infection from individuals already infected at time 0 is instead captured in the new term 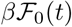, where

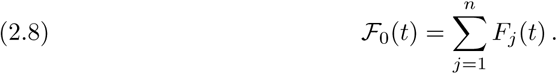

The *F_j_*’s are calculated by integrating the cohort equations (2.3) starting from the desired initial conditions, which can be done in advance (either analytically or numerically) or simultaneous with numerically solving the alternative form of the RE (Equations 2.5a and 2.7).

## 3. Results

### 3.1. The intrinsic generation-interval distribution of the Erlang SEIR model

Here, we solve the ODE system (2.3) in order to obtain an analytical expression for the generation-interval distribution *g* for an Erlang SEIR model, using Equation 2.4.

Because Equation 2.3 is a linear ODE, it can be solved exactly. Solving for the probabilities to be in the *j*^th^ latent stage *L_j_* is straightforward. Equation (2.3a) gives *L*_1_(*t*) = *e*^−(^*^mσ^*^+^*^µ^*^)^*^t^*. Multiplying equation (2.3b) by *e*^(^*^mσ^*^+^*^µ^*^)^*^t^* for *k* = 2 gives (*e*^(^*^mσ^*^+^*^µ^*^)^*^t^L*2)′ = *mσ*, hence *L*_2_(*t*) = *mσte*^−(^*^mσ^*^+^*^µ^*^)^*^t^* (recall that *L*_2_(0) = 0). It is then easy to prove by induction that

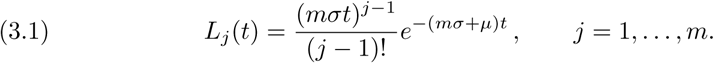

Solving for the probabilities to be in the *k*^th^ infectious stage *F_k_* is more tedious. We present the two special cases when *m* = 0 and *mσ* = *nγ* first because both the calculations and expressions are much simpler, then we give the expression for the general case.

#### 3.1.1. Case *m* = 0

If *m* = 0 (which is also equivalent to *σ* → ∞), then the *F_k_* satisfy the same ODE as the *L_k_* in the case where *m* > 1. Hence, we have:

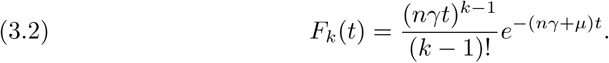

The integration is straightforward:

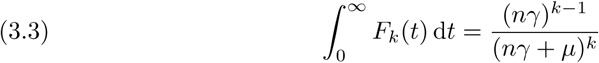

Using Equation 2.4, the intrinsic generation-interval distribution is

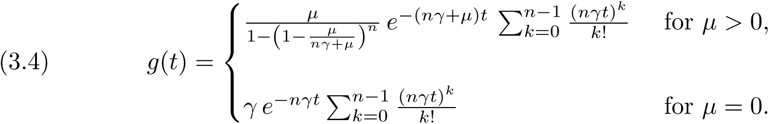

In the special case *n* = 1 this reduces to

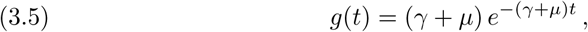
recovering the well-known result that the standard SIR model has an exponential intrinsic generation-interval distribution [4].

#### 3.1.2. Case *m* > 0 but *mσ* = *nγ*

If *mσ* = *nγ*, the analytical expression for *F_k_* is obtained in a similar way as *L_k_*:

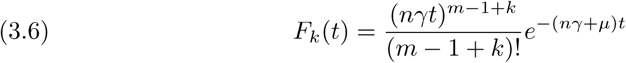

The integration is again straightforward and we have

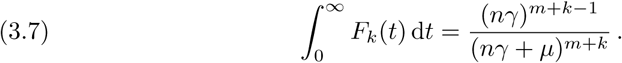

Hence, using Equation 2.4 the intrinsic generation-interval distribution is

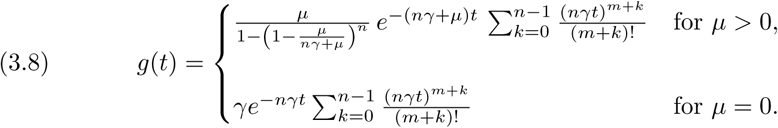

In the special case of the standard SEIR model (*m* = *n* = 1), for any *µ* ≥ 0, we obtain

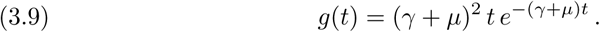

#### 3.1.3. General case *m* > 0 and *mσ* ≠ *nγ*

In this case, we set *µ* = 0 as it simplifies both the calculations and expressions considerably. For typical epidemics of infectious disease, the demographic rate *µ* is usually negligible compared to the epidemiological rates (i.e., *µ* ≪ *mσ* and *µ* ≪ *nγ*), so the effect of *µ* on the generation interval distribution *g*(*τ*) will also be negligible in most applications. Calculations described in Appendix A yield

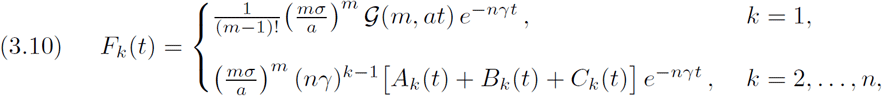

where

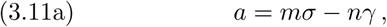

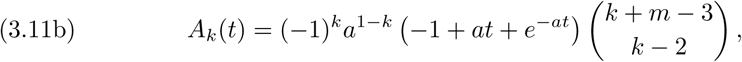

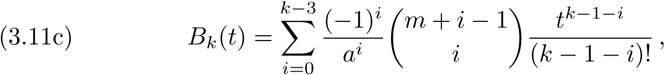

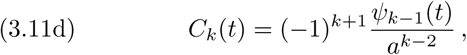

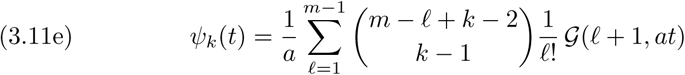

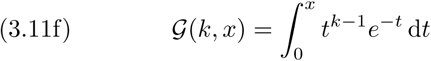

The function 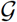 is the lower incomplete gamma function [23, §8.2.1]. We obtain the intrinsic generation-interval distribution for the Erlang SEIR by combining equations (2.4) and (3.10). In this generic case we obtain

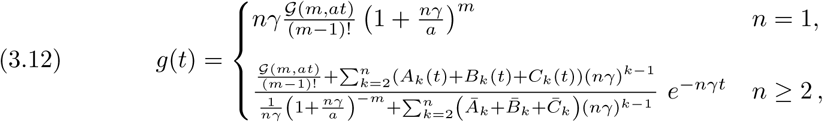

where

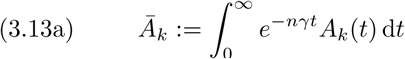

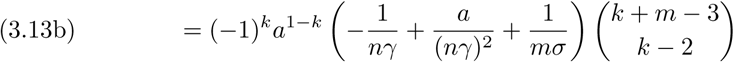

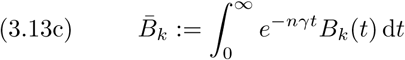

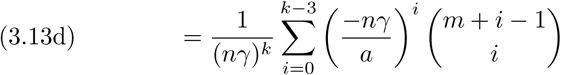

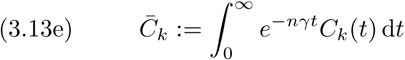

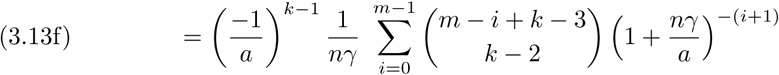

In the special case *m* = *n* = 1, i.e., the standard SEIR model, all the complexities collapse and we obtain

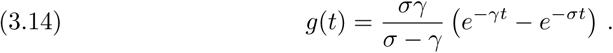

#### 3.1.4. Discrete time SIR

While our focus has been on continuous-time models, it is worth mentioning that the SIR model in discrete time is equivalent to the renewal equation with a geometric generation-interval distribution, with probability parameter *γ*Δ*t*, where Δ*t* is the time discretization step (which must be chosen such that *γ*Δ*t* < 1). This result, which we derive in Appendix B, is consistent with the fact that the exponential distribution is the continuous analog of the geometric distribution.

### 3.2. Numerical simulations

We verified the correctness of our analytical expressions for the stage duration distributions, equations (3.1) and (3.10), by comparing them with direct numerical integration of the linear ODE system for these probabilities (2.3). Figure A1 shows a visually perfect match between the analytical formulae and the numerical solutions for *L_k_*(*τ*) and *F_k_*(*τ*). Inserting Equation 3.10 into Equation 2.4 we obtained the associated intrinsic generation-interval distribution *g*(*τ*), which is plotted in Figure A2 together with the approximate distribution obtained by integrating the linear ODEs (2.3) numerically.

We then checked that solutions of the renewal equation (2.5) agree with those of the Erlang SEIR ODE system (2.1). As an example, Figure 1 shows a visually perfect match between the two models for a particular parameter set.

**Fig. 1.**
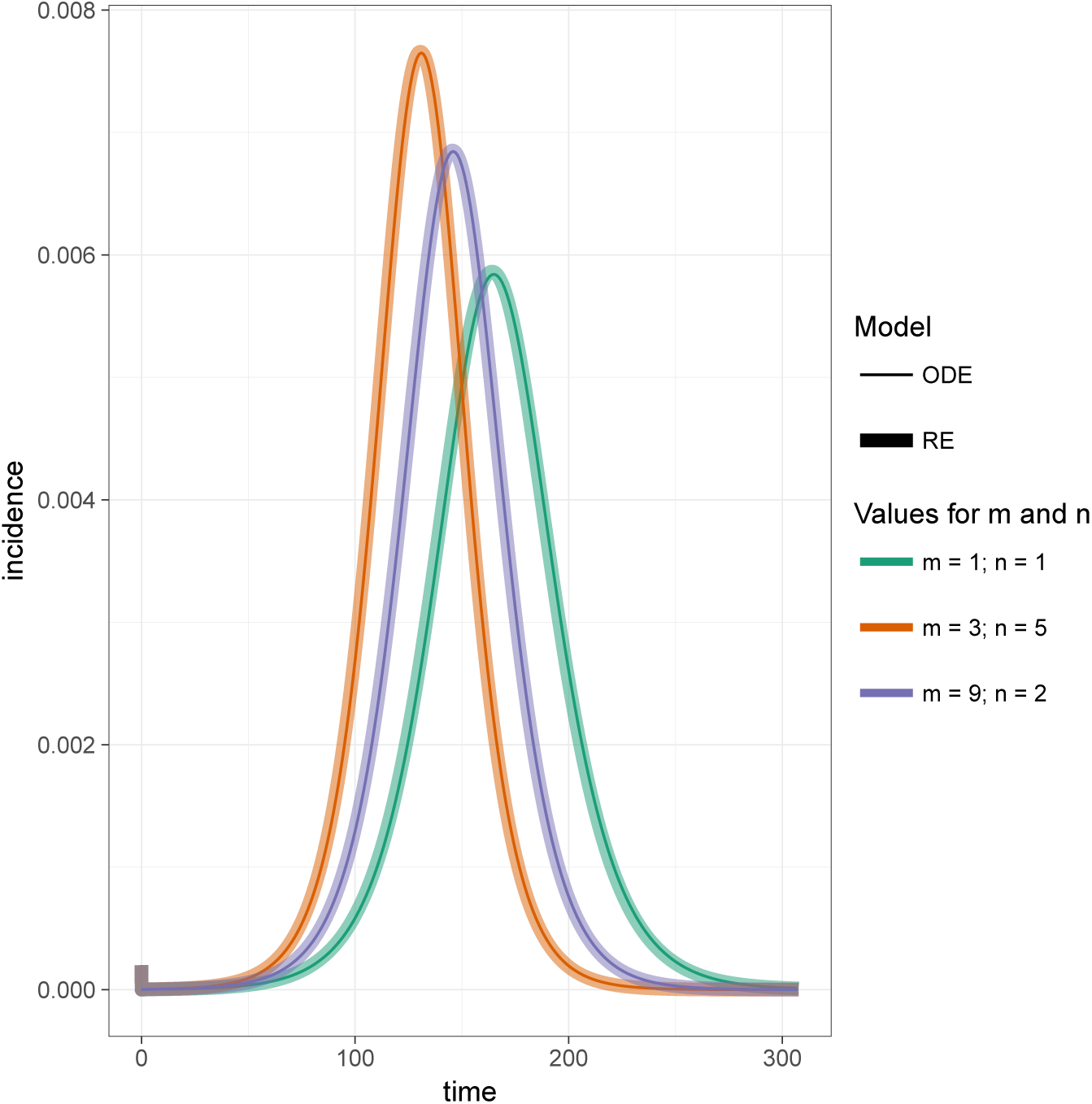
Numerical check of equivalence in continuous time. *Daily incidence time series of the Erlang SEIR for different values of m and n is obtained by solving numerically the ODE system (2.1) (and retrieve βSI as the incidence). The daily incidence time series of the renewal equation (RE) was calculated using equation (2.5) and C.1 with the intrinsic generation interval* g *defined with formula (3.12) and a time step of 0.1 day. The superimposed curves (solid line for ODE and dash for RE) show the equivalence of both models when the generation-interval distribution of the renewal equation is appropriately chosen. Mean duration of latency (respectively infectiousness) is 2 (respectively 3) days, and* 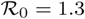.

We also checked our finding that the discrete time SIR model (§3.1.4 and Appendix B) is equivalent to a renewal equation model with a geometric generation interval distribution (Figure B1). Moreover, Figure 2 shows an illustrative example of the equivalence of the renewal equation (2.5) and the Erlang ODE system (2.1) in the presence of vital dynamics and periodic forcing of the transmission rate. In this example we used again the RE model with a geometric generation interval distribution, and applied a sinusoidally forced basic reproduction number 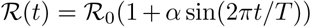 with 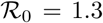, forcing amplitude *α* = 0.6, forcing period *T* = 365 days, and birth and death rates *µ* = 0.03 yr^−1^.

**Fig. 2.**
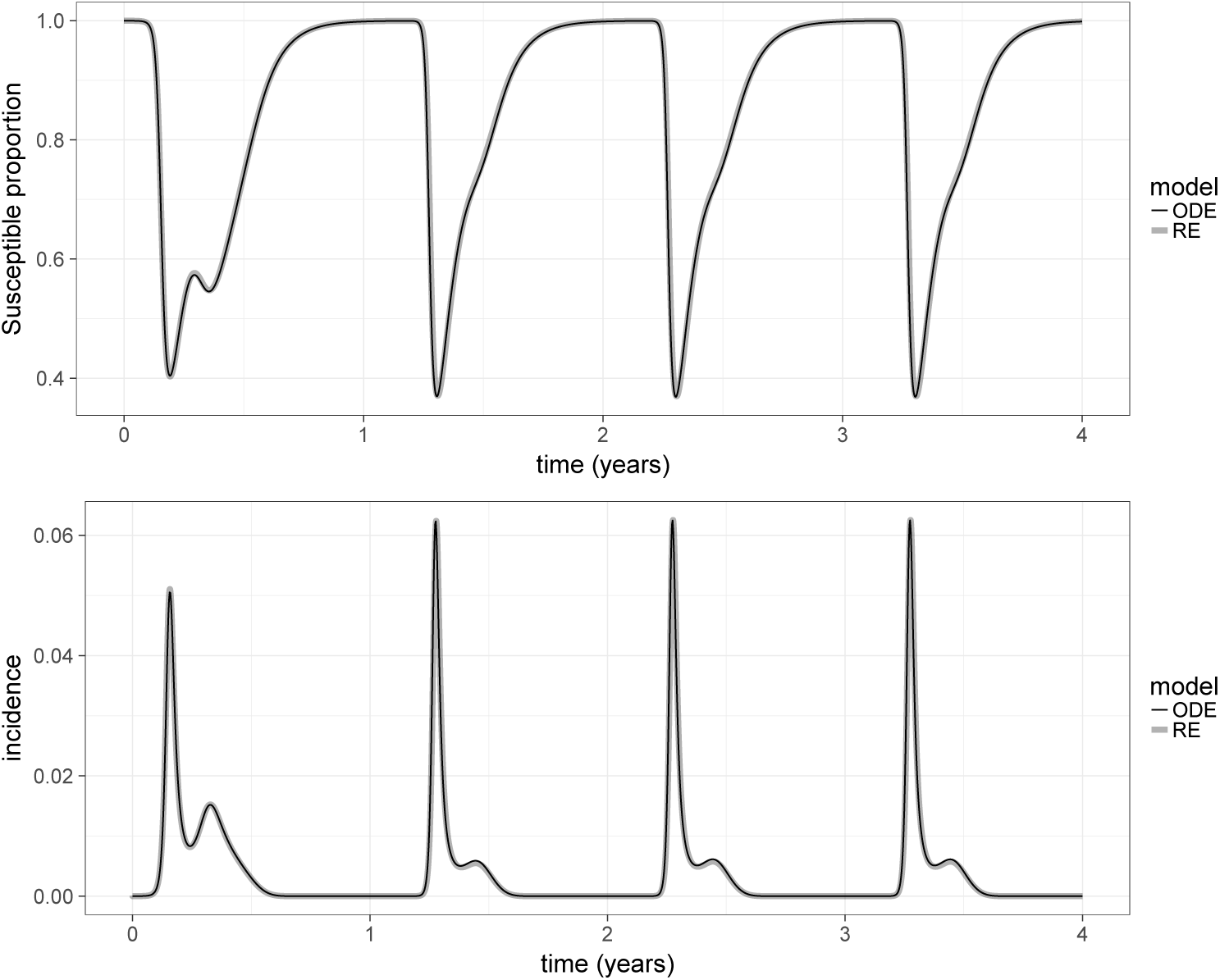
Time series for a SIR model with vital dynamics and seasonal forcing. *Top panel: susceptible proportion; bottom panel: daily incidence. The thin black curve represents the time series obtained by solving numerically the ODE system (2.1). The thick grey time series was calculated using the renewal equation model (2.5) with an exponential intrinsic generation-interval distribution and implemented with an integration time step of 0.05 day. The birth and death rate is µ* = 0.02/*year, the mean infectious period is* 1/*γ* = 3 *days. The reproduction number was periodically forced* 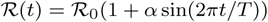 *with* 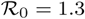, *α* = 0.6 *and T* = 365 *days*.

## 4. Discussion

Appreciation of the the fact that many epidemic models can be expressed either with ordinary differential equations (ODEs) or with a renewal equation (RE) can be traced back to the original landmark paper of Kermack and McKendrick [17, 5]. Provided one wishes to track only the dynamics of the total susceptible population and incidence rate, there is no difference in the output of the two formulations. This result is well known in the broader field of delayed integro-differential equations [11, 28] (and sometimes described as the “linear chain trick” [5]). While references to this connection have certainly been made in epidemiological contexts (see for example [12, 15, 5]), the epidemic modelling community has not taken full advantage of this result. Here, by providing exact analytical expressions for the intrinsic generation-interval distribution of any Erlang SEIR model, we hope to draw attention to the renewal equation and its potential uses in studying infectious disease dynamics. Table 1 summarizes our main results. We note that the methodology we have used to derive the intrinsic generation interval distribution *g*(*τ*) required in the renewal equation (2.5) can be applied to any staged-progression epidemic model [16].

Epidemic models described by ODEs—with state variables corresponding to compartments that represent various epidemiological states—are invaluable tools for evaluating public health strategies [2]. For example, when the goal of a modelling study is to assess a particular intervention (e.g., vaccination of a particular group) in a large population, a compartmental ODE is convenient because it is easy to keep track of the numbers of individuals in each disease state. The Erlang SEIR model is often a good choice, at least as a starting point, because it can represent realistic distributions of latent and infectious periods [26]. However, if one is interested only in the dynamics of the susceptible and/or infectious populations (e.g., when forecasting incidence in real time during an outbreak), the renewal equation framework can be beneficial as it can simplify the modelling [8] and potentially speed up the computation times. The analytical formulae for the intrinsic generation interval of the SEIR Erlang ODE model (equations (3.4), (3.8), (3.12), or Table 1) are relatively easy to implement in a computer program. Our experience has been that the renewal equation yields faster numerical simulations than the corresponding ODE models. Of course, computing times depend on the numerical methods and software implementation; more work is needed to ascertain how computing times vary between approaches given identical problems and equivalent error bounds.

The generation interval is rarely observed, but through contact tracing it is possible to directly observe the *serial interval* (i.e., the interval of time between onset of symptoms for the infector and her/his infectee). Although different in theory, the serial interval distribution may be a good approximation to the generation interval distribution, especially for diseases for which the latent and *incubation* periods are similar (Appendix D and [14]). On the other hand, the latent and *infectious* periods—which are used to parametrize compartmental ODE models—can be observed only in clinical studies, which are more rare. Consequently, the generation-interval distribution can be easier to obtain than the distributions of latent and infectious periods, in which case a renewal equation might be easier to parameterize than an Erlang SEIR ODE model.

**Table 1.** *Compartmental models and their equivalent intrinsic generation interval distribution for the renewal equation. The mean duration of the latent (resp. infectious) period is* 1/*σ (resp*. 1/*γ). The variable t is the time since infection and* Δ*t (which must be less than* 1/*γ) is the size of the time step when time is discrete. If µ* > 0 *then one just replaces σ and γ with σ* + *µ and γ* + *µ in g*(*τ*) *for SIR and SEIR*.

| Compartmental<br>ODE | RE intrinsic generation-interval distribution $g(t)$ |
| --- | --- |
| $SIR$ discrete time | Geometric($\gamma\Delta t$): $\gamma\Delta t(1 - \gamma\Delta t)^{\frac{t}{\Delta t}-1}$ |
| $SIR$ | Exponential($\gamma$): $\gamma e^{-\gamma t}$ |
| $SEIR$ | $\begin{cases} \gamma^2 t e^{-\gamma t} & \sigma = \gamma \\ \frac{\sigma\gamma}{\sigma-\gamma} (e^{-\gamma t} - e^{-\sigma t}) & \sigma \neq \gamma \end{cases}$ |
| $SE^m I^n R$ (“Erlang”) | $\begin{cases} \text{Equation 3.4} & m = 0 \\ \text{Equation 3.8} & m\sigma = n\gamma, m > 0 \\ \text{Equation 3.12} & m\sigma \neq n\gamma, m > 0 \end{cases}$ |

## Appendix

## Appendix A. Proof of formula (3.10)

### A.1. Preliminaries

**Lower incomplete gamma function**. The lower incomplete gamma function 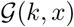 is defined for *k* > 0 and *x* ≥ 0 via [23, §8.2.1]

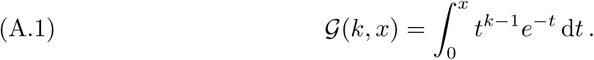

We use the notation 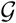 rather than the standard *γ* for this function because, in this paper, we reserve the symbol *γ* for the disease recovery rate. The integral of 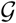 can be written

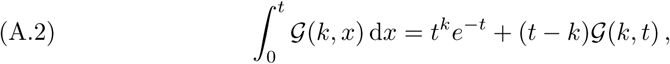
which is straightforward to verify by noting that both sides vanish for *t* = 0 and that they have identical derivatives. Because it is an expression that occurs often in our calculations, we note that

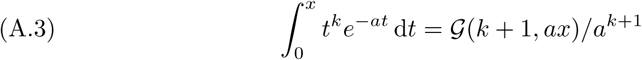

**Nested sums**. In the course of our computations, certain types of nested sums occur repeatedly, so it is helpful to note that, for any function *f*,

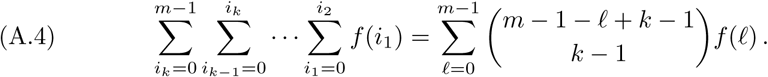

In the special case *f*(0) = 0 and *f*(*ℓ*) = 1 for all *ℓ* ≥ 1, we have [6]

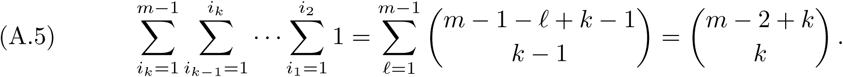

We define for any integers *m* > 0, *k* > 0 and real *a*,

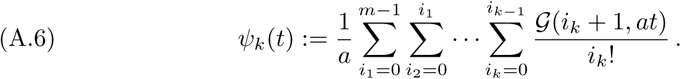

Using Equation A.4, we can re-write *ψ* as a single sum,

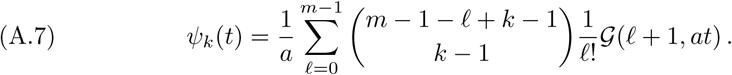

It can then be proved by induction that the integral of *ψ_k_* is

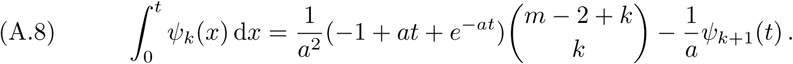

We note, in particular, that *ψ*_0_ = 0 and 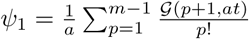.

### A.2. Calculations for *F*_1_

We first consider *F*_1_. From the system of ODEs Equation 2.3 (in the main text) we have:

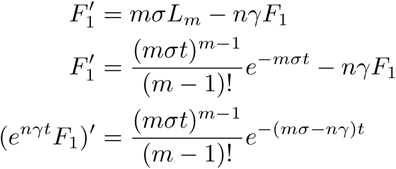

Hence,

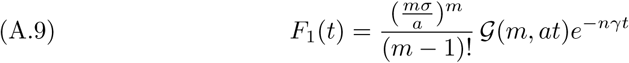

### A.3. Calculations for *F_k_* for *k* ≥ 2

Again from Equation 2.3 we have 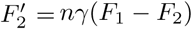. Multiplying both sides by *e^nγt^* gives

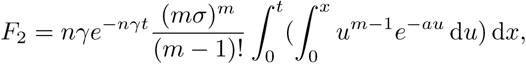
which can be expressed explicitly using the lower incomplete gamma function,

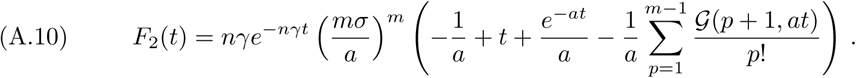

Similarly, starting from 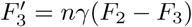 and multiplying both sides by *e^nγt^* we have, after some algebra,

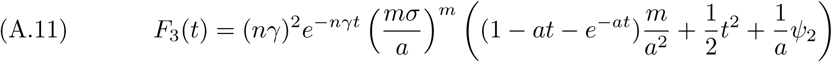

Using the results from subsection A.1, we can prove by induction (using *F*_3_ as the initial step) that

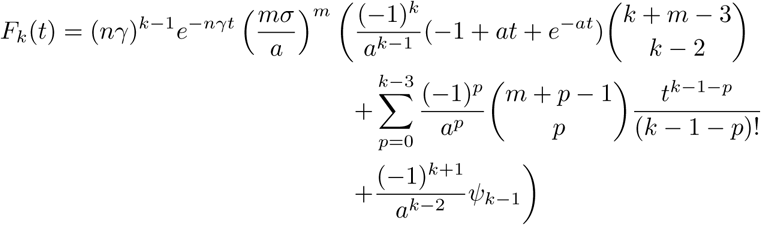

**Fig. A1.**
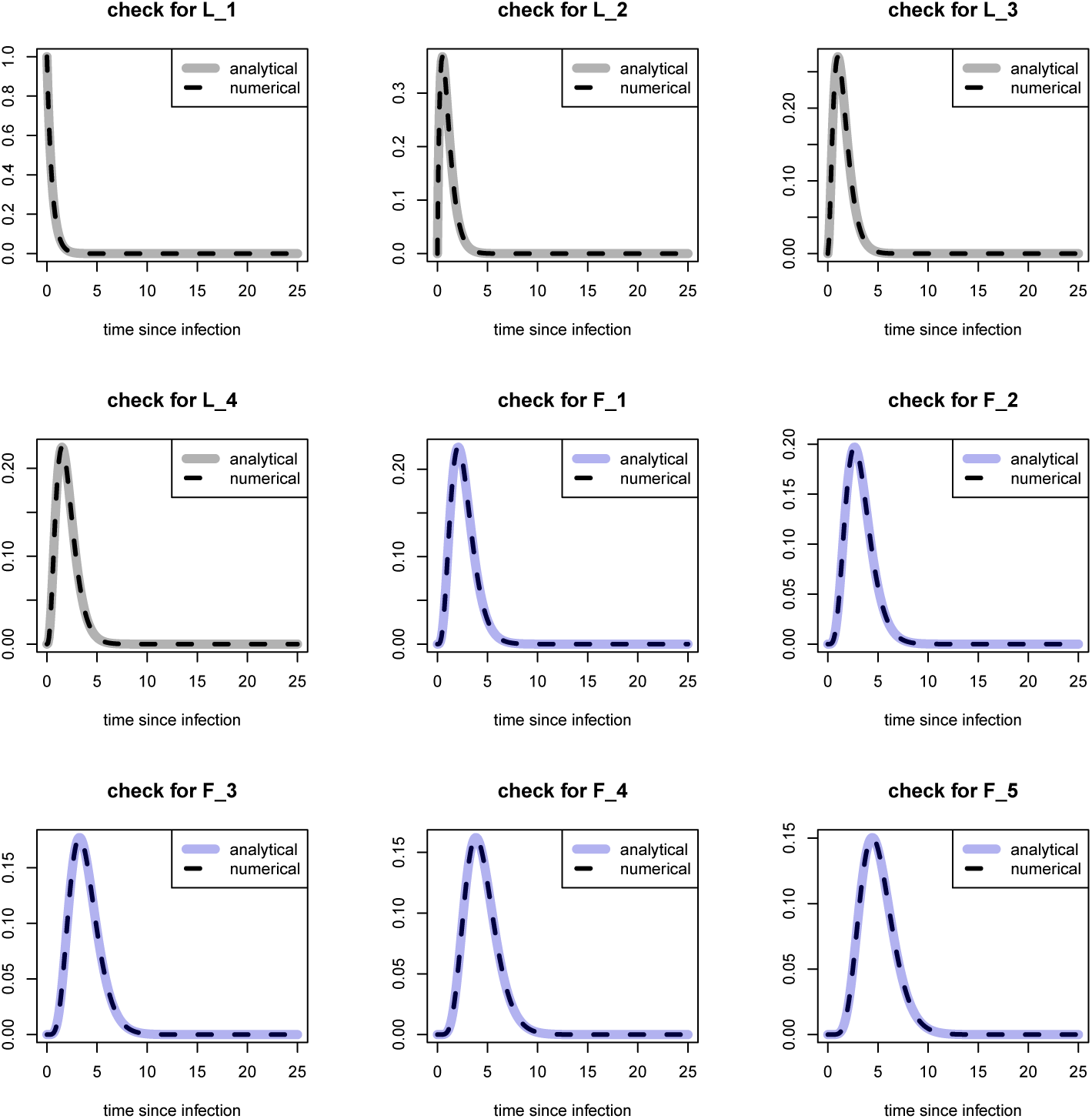
*Formula checks for probabilities L_k_ and F_k_. Analytical formulas (3.1) and (3.10) are compared to the numerical integration of the ODE system (2.3). For this figure, m* = 4 *and n* = 5.

## Appendix B. Renewal equation and SIR model in discrete time

This section has the pedagogical purpose to show how, in the case of a discrete time SIR model, the generation interval distribution can be calculated using simple mathematical manipulations.

### B.1. Discrete time

The discrete time formulation of the renewal equation, without vital dynamics, is:

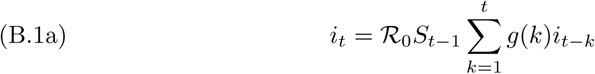

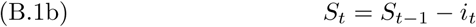

The SIR model has been extensively covered. We consider a standard, discretized version of an SIR model without vital dynamics, where the incidence *i_t_* is introduced explicitly:

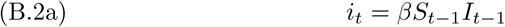

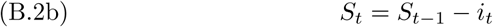

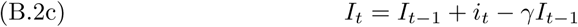

**Fig. A2.**
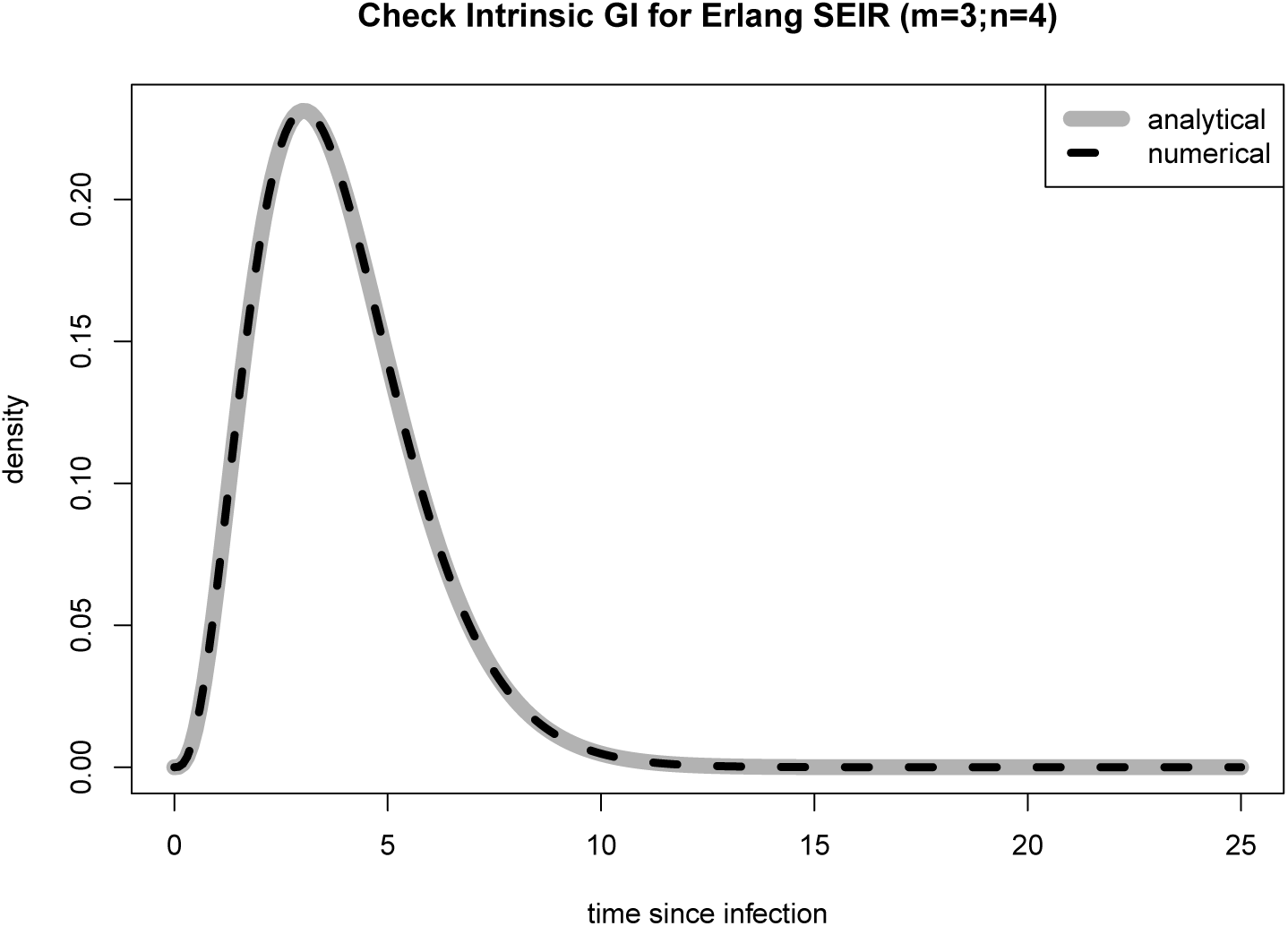
*Formula check for the intrinsic generation-interval distribution. Analytical formulas (2.4) and (3.10) are compared to the numerical integration of the ODE system (2.3) when m* = 3 *and n* = 4.

When studying disease invasion, we take initial conditions *I*_0_ = 1 − *S*_0_ ≪ 1. We note that equation (B.2c) can be rewritten as *I_t_* = (1 − *γ*)*I_t_*_−1_ + *i_t_*. Substituting *I_t_*_−1_ = (1 − *γ*)*I_t_*_−2_ + *i_t_*_−1_ gives: *I_t_* = (1 − *γ*)^2^*I_t_*_−2_ + (1 − *γ*)*i_t_*_−1_ + *i_t_*. Iterating this substitution *t* times, we have:

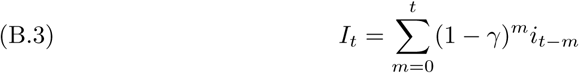

Next, we use equation (B.2a) to substitute the left hand side by *I_t_* = *i_t_*_+1_/*βS_t_*, operate a shift of one time unit *t* → *t* − 1:

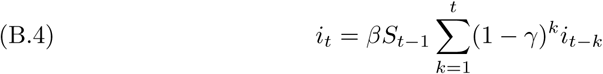

If we note 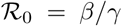, set 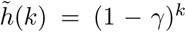 and the normalized function 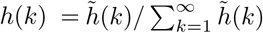 we have:

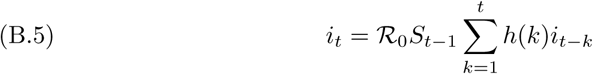

Hence, we have expressed the SIR model in the same form as the renewal equation. The function *h* can then be identified as the intrinsic generation-interval distribution in the renewal equation framework. We have *h*(*k*) = *γ*(1 − *γ*)*^k^*^−1^, which is the density of the geometric distribution with probability parameter *γ*. Hence, a discretized SIR model is exactly the same as a renewal equation model with a geometric generation-interval distribution.

### B.2. Limit of continuous time

We will also need an expression of the renewal equation when using a time step that is smaller than the time unit (i.e., day). The renewal equation models how transmission occurs from all previous cohorts (infected at times 0, 1, …, *t* − 1) to the current time (*t*). The way the generation-interval distribution *g* is defined depends on the unit of the time discretization. Writing the renewal equation (B.1a) necessitates changing the definition of incidence from daily incidence to incidence during the new time step period. Moreover, if we want to keep the same parameterization for the generation-interval distribution, then *γ* must be rescaled. Let’s consider a time step *δ* < 1 that partitions time in *N* segments of the same size (*δ* = 1/*N*). Rewriting the renewal equation (B.1a) with that new subpartition gives, for any *m* > 0:

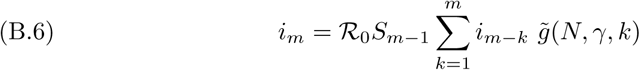

Despite using the same notations, the implicit meaning for *i* and *S* in Equation (B.6) has changed and now refers to the incidence and susceptible proportion during the time step Δ*t* (not 1 day). The index *k* now refers to new *k*^th^ period of length Δ*t*. Moreover, the generation-interval distribution 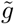 now takes into account the time scale change, while keeping the same parameterization with *γ*. In Equation (B.1a), taking a geometric distribution for the generation interval, *g*(*k*)= *γ*(1 − *γ*)*^k^*, so the mean generation interval is 1/*γ* in the original time unit (e.g., days). If we were to write *g*(*m*) = *p*(1−*p*)*^m^* in Equation (B.6), the mean generation interval would be 1/*p* in the new time unit (e.g., hours). Hence we must have 1/*p* = *N* × 1/*γ*, that is *p* = *γ*/*N*. So, we have 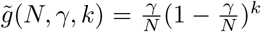. To summarize, the renewal equation written for a time step Δ*t* = 1/*N* of the original (natural) time, assuming a geometric distribution with mean 1/*γ* original time unit for the generation interval is

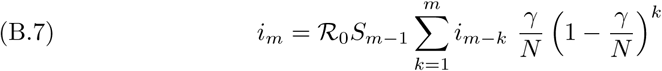

Now we consider an arbitrary small subpartition of the (natural) discrete time and will take the limit when the time step tends to 0 in order to have results in continuous time.

Starting again with the SIR model for time step of Δ*t* = 1/*N*, we can rewrite Equation (B.2c) as (*I_k_* − *I_k_*_−1_)/(1/*N*) = *i_k_* − *γI_k_*_−1_ that is *I_k_* = *i_k_* − (1 − *γ*/*N*)*I_k_*_−1_, where *I_k_* and *i_k_* now refer to the prevalence and incidence of the *k^th^* period of length Δ*t*. Using the same algebraic manipulations as in the previous section with the original time unit, gives the following expression for the incidence during the *m*^th^ period of an SIR model:

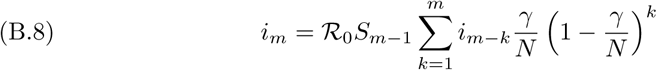
which is exactly the same as the renewal equation (B.7). Hence, the result obtained for the original (natural) time discretization—i.e., the discretized renewal equation with a geometric generation interval is the same as the discretized SIR model—still holds for any subpartitioned time discretization, as long as the probability parameter of the geometric distribution is rescaled accordingly (i.e., 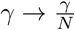).

For both the SIR model and the renewal equation, the continuous time formulation is obtained when taking the limit *N* → +∞ (that is, Δ*t* → 0). But the limit of the geometric distribution in equation (B.7) is the exponential distribution. Hence, the continuous time formulation of the SIR model is equivalent to the continuous time formulation of the renewal equation with an exponential distribution for its generation interval.

**Fig. B1.**
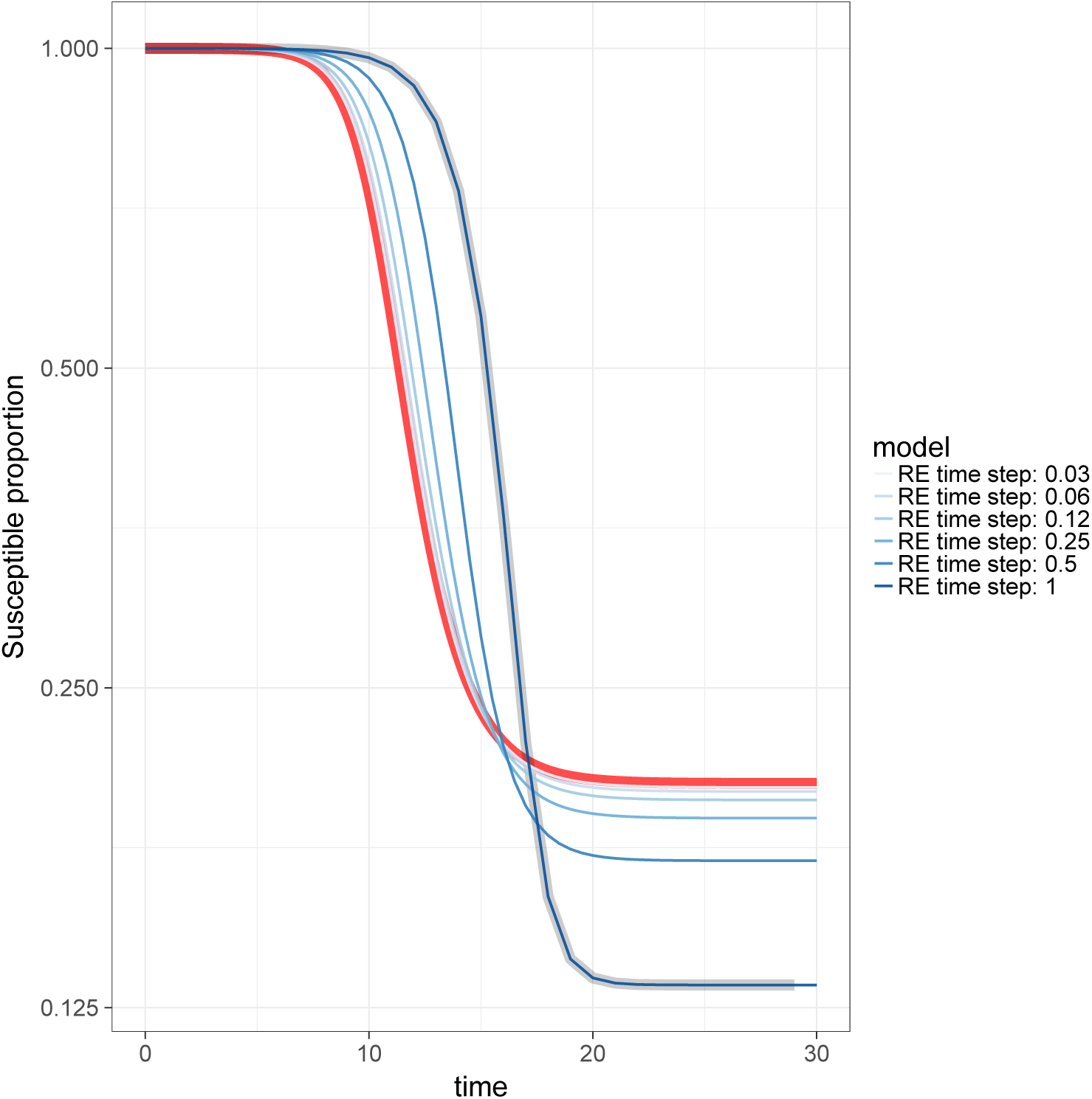
Numerical check of equivalence in discrete time. *The thick lines show the time series of the susceptible proportion of the population for the SIR model in discrete time (time step = 1, grey curve) and in continuous time (time step = 0.01, red). The thin blue lines represent the susceptible proportion from the discrete renewal equation (RE) implemented with different time step values* Δ*t* = 1/*N with N* = 1, 2, 4, 8, 16, 32*. The RE model has a generation interval geometrically distributed, with the time-rescaled probability parameter γ*Δ*t (Equation B.7). When N* = 1 *the RE is simulated at the same times as the discrete SIR, and the two curves match. As N increases, time discretization becomes closer to continuous time and the RE curves approach the SIR model simulated in continuous time. The y-axis has a log scale to better visualize the difference between the curves. Parameters used:* 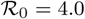*, mean duration of infection γ* = 1 day^−1^*, initial proportion of infectious individuals I*_0_ = 10^−5^.

## Appendix C. Numerical solution of the renewal equation

The Rewewal Equation 2.5 with invasion initial conditions (2.6) can be solved, for an integration time step Δ*t*, using the “left Riemann sum” approach detailed in C.1.

#### Algorithm C.1 Numerical simulation of the renewal equation

**Input**: Positive real number 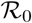, *µ*, and *t*_max_; initial prevalence *I*_0_; density function *g*; integration time step Δ*t*

\## initialization

inc[0] ← *I*_0_/Δ*t*

*S*[0] ← 1 *− I*_0_

nsteps ← *t*_max_/Δ*t*

\## Loop calculating incidence at each time step

integ ← 0

**for** (*u*=1, 2, …, nsteps) **do**

    integ ← integ + *g*(*s*) * inc[*u* − *s*] * exp(*−µ* * Δ*t* * *s*)

    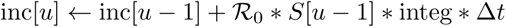

    *S*[*u*] ← *S*[*u* − 1] + (*µ* * (1 − *S*[*u* − 1]) − inc[*u*]) * Δ*t*

**end for**

**return** Proportion of susceptible (vector *S*) and incidence (vector inc).

## Appendix D. Generation and serial intervals distributions

As highlighted in [14], there are three fundamental time periods that determine transmission from one individual to another for directly-transmitted infectious diseases: the latent, incubation and infectiousness periods. Let *ℓ*_1_ be the latent period of an infector and *ℓ*_2_ the latent period of her/his infectee. Let *w* the interval of time between the end of the infector’s latent period and the time of disease transmission to an infectee. We note *n*_1_ and *n*_2_ the incubation period of the infector and infectee, respectively. The difference between the latent and incubation periods is noted *d_i_* = *ℓ_i_* − *n_i_* for *i* = 1, 2. The generation interval is *g* = *ℓ*_1_ + *w* and the serial interval is *s* = (*ℓ*_1_ + *w* − *n*_1_) + *n*_2_ (Figure D1). Hence we can write

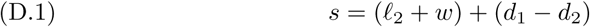

**Fig. D1.**
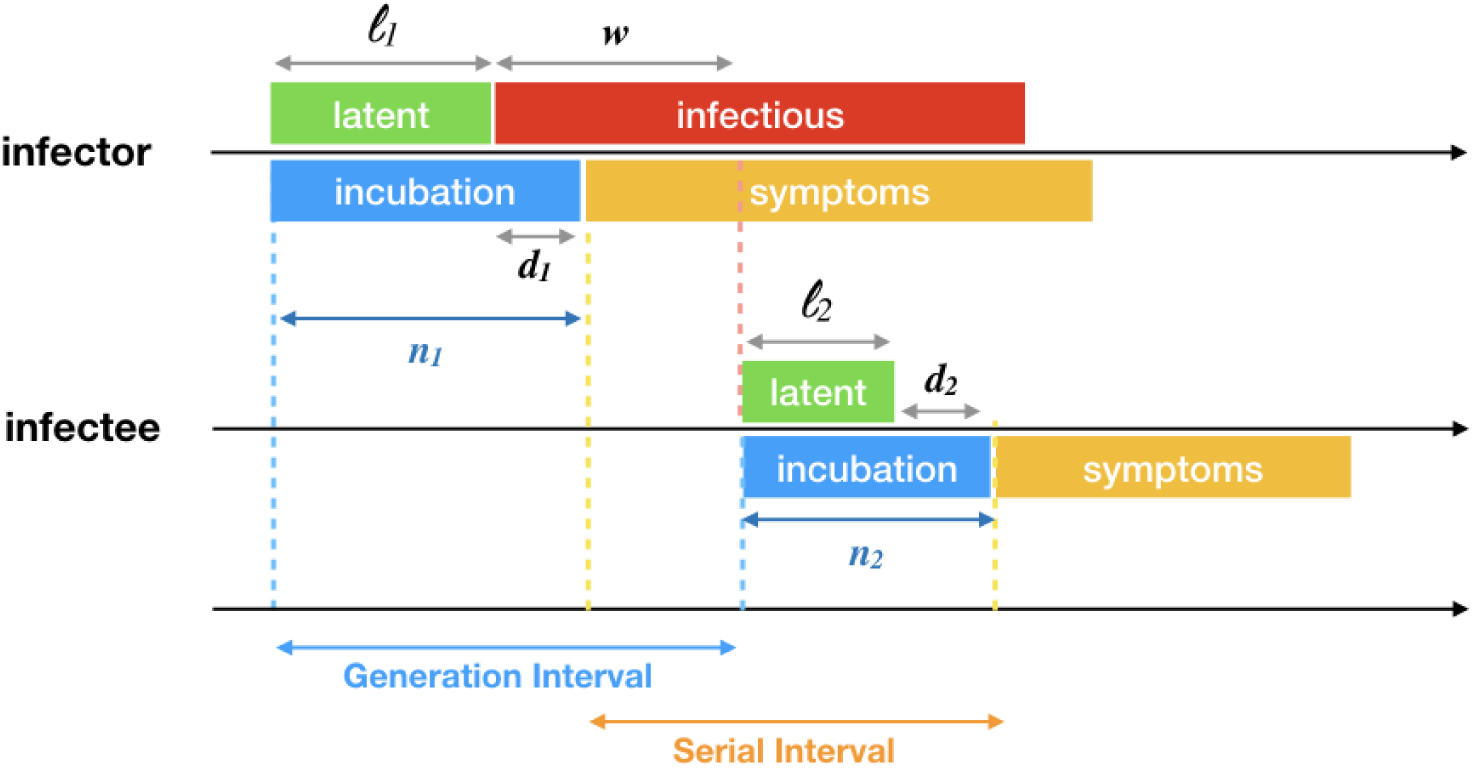
*Illustration of the epidemiological periods and parameters*.

If we assume that *ℓ*_1_ and *ℓ*_2_ are identically distributed, and also *d*_1_ and *d*_2_ are identically distributed with distribution 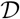, then the generation interval distribution 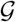 and serial interval distribution 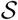 have the same mean:

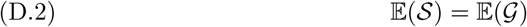

If furthermore we assume that *d*_1_ and *d*_2_ are independent from one another, and also from *ℓ* and *w*, we can write:

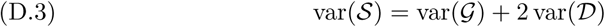

So when the variance of the difference between the latent and incubation periods is small, the variance of the serial and generation intervals are similar.

## Footnotes

* Submitted to the editors May 10, 2018 at 12:14:27.

1 Or, more generally, susceptible recruitment.

